# Inductance in Neural Systems

**DOI:** 10.1101/343905

**Authors:** Hao Wang, Jiahui Wang, Xin Yuan Thow, Sanghoon Lee, Wendy Yen Xian Peh, Kian Ann Ng, Tianyiyi He, Nitish V. Thakor, Chia-Hung Chen, Chengkuo Lee

## Abstract

A neural circuit model involving inductance is established to explain the neural networks’ behavior. A parallel resistor-inductor-capacitor (RLC) circuit was used to fit the stimulus artifacts in the electromyography (EMG) recording of cortical and pelvic electrical nerve stimulations. This parallel RLC circuit model also predicts the resonance effect in both stimulus artifacts and EMG signals. Moreover, the well-known strength-duration relationship was directly derived to be a precise format with this parallel RLC circuit model. A theoretical explanation is provided to show the inductance is generated by the coil structure of the myelin sheath and the piezoelectric effect of the plasma membrane.

**One Sentence Summary:** The inductance in the neural systems is generated by the coil structure of the myelin sheath and the piezoelectric effect of the plasma membrane.

## Main Text

Neural circuit model is a popular approach and a useful mathematical model to investigate the electrical current and neural signal propagation on <u>neurites</u> at different sites and times *(1)*. A most critical issue for this approach is the configuration of the circuit. Starting from the cable model, which is directly derived from the biological structure of the axon, and developed by the famous Hodgkin–Huxley model (H-H model) *(2)*, the RC (resistor-capacitor) configuration has been a basic circuitry unit for whole computational neuroscience. The existence of the inductance has never drawn the attention of the mainstream academia. However, even in the early stage of neuroscience, the direct impedance measurement of giant squid axon already confirms the inductive reactance in axons *(3)*. This huge inductance can be repeatedly measured in all kinds of neuron related research *(4-18)*. Moreover, some abnormal experimental observations, such as the frequency dependent response of nerve fibers *(19-23)* and the anode break excitation *(24,25)*, cannot be well explained without including the inductor in the neural circuit. To solve these issues, some theories and ideas are proposed, such as frequency-dependent membrane capacitance *(26)* (for the frequency dependent response) and the virtual-cathode hypothesis *(27)* (for the anode break excitation). To develop a comprehensive model, an inductor was added in the neural circuit. With the inductor, the neural circuit should follow an RLC configuration rather than an RC configuration. Then the frequency dependent response becomes the intrinsic feature of the neural system. The anode break excitation is also comprehensible: the biphasic voltage oscillation of the RLC circuit after the termination of a <u>hyperpolarizing</u> current, which will be detailed demonstrated in this study, surely has a chance to fire the action potential. Is there an inductance factor in the neural systems? The answer is “Yes”. Previously people tend to believe this inductance does not really exist simply because they cannot find a physical entity to account for such a huge inductive reactance. In this study, together with the direct experimental evidence of the existence of the inductance in neural systems, a comprehensive theoretical explanation of the physical entities to generate this inductance will be provided. To proof the correctness of this theory, we will show how those most important experimental observations and phenomena are derived from this theory.

In the experiment part, we analyzed the signal in EMG recording of electrical nerve stimulations. The curve fitting of the stimulus artifact shows that the neural circuit should generally follow a parallel RLC configuration. The resonance frequency of this RLC circuit can be measured in both stimulus artifacts and EMG signals. Then based on this RLC circuit, the well-known strength-duration relationship *(28,29)*, can be directly derived from this parallel RLC circuit. Moreover, this strength-duration relationship is amended to a more precise format and its prediction has already been validated by experimental observations from other groups *(30-32)*.

In the theory part, a comprehensive theoretical explanation of the physical entities to generate the inductance is provided. Simply speaking, this inductance is generated by the coil structure of the myelin sheaths and the piezoelectric effect of the plasma membrane. Based on this theory, we can get many unique predictions, which are all validated by previous researches and observations. Here we only give the two most critical predictions:

1. Adjacent myelin sheaths always follow the opposite wrapping orientations, which has been well validated by the previous study *(33-35)*. This phenomenon cannot be explained by any previous theories and models, but now can be obtained from this theory.
2. There will be a mechanical wave accompany with the electrical neural signal. This mechanical wave has been studied for more than 10 years and an alternative theory called soliton theory has been developed to explain this mechanical wave *(36)*. In soliton theory, with a complicated calculation, it is claimed that the propagation speed of the mechanical wave is the same as that of the electrical signal predicted by the H-H model. However, in our theory, we can get this result without any calculation. This mechanical wave is an attendant phenomenon with the electric neural signal. It surely has the same speed as the electric signal.

## Experimental data showing the evidence of the inductance in neural systems

### A. Analysis of the stimulus artifacts

The typical samples of the EMG recording in the electrical stimulation of the cortical and pelvic nerves are shown in Figure 1(a) and (c), respectively. The experimental details can be found in **Supplementary S1 and S2**. As seen, there will be two separate signals: the stimulus artifact and the EMG signal induced by the electrical stimulation. For most of the cases in previous studies *(37,38)*, people only focused on the analysis of the EMG signals since they are the real neural responses. However, the stimulus artifacts are the direct electrical responses of the whole tissue and organism affected by the injected current, revealing its electrical characteristics. In other words, the configuration of the equivalent neural circuit can be obtained from these electrical responses.

**Fig. 1.**
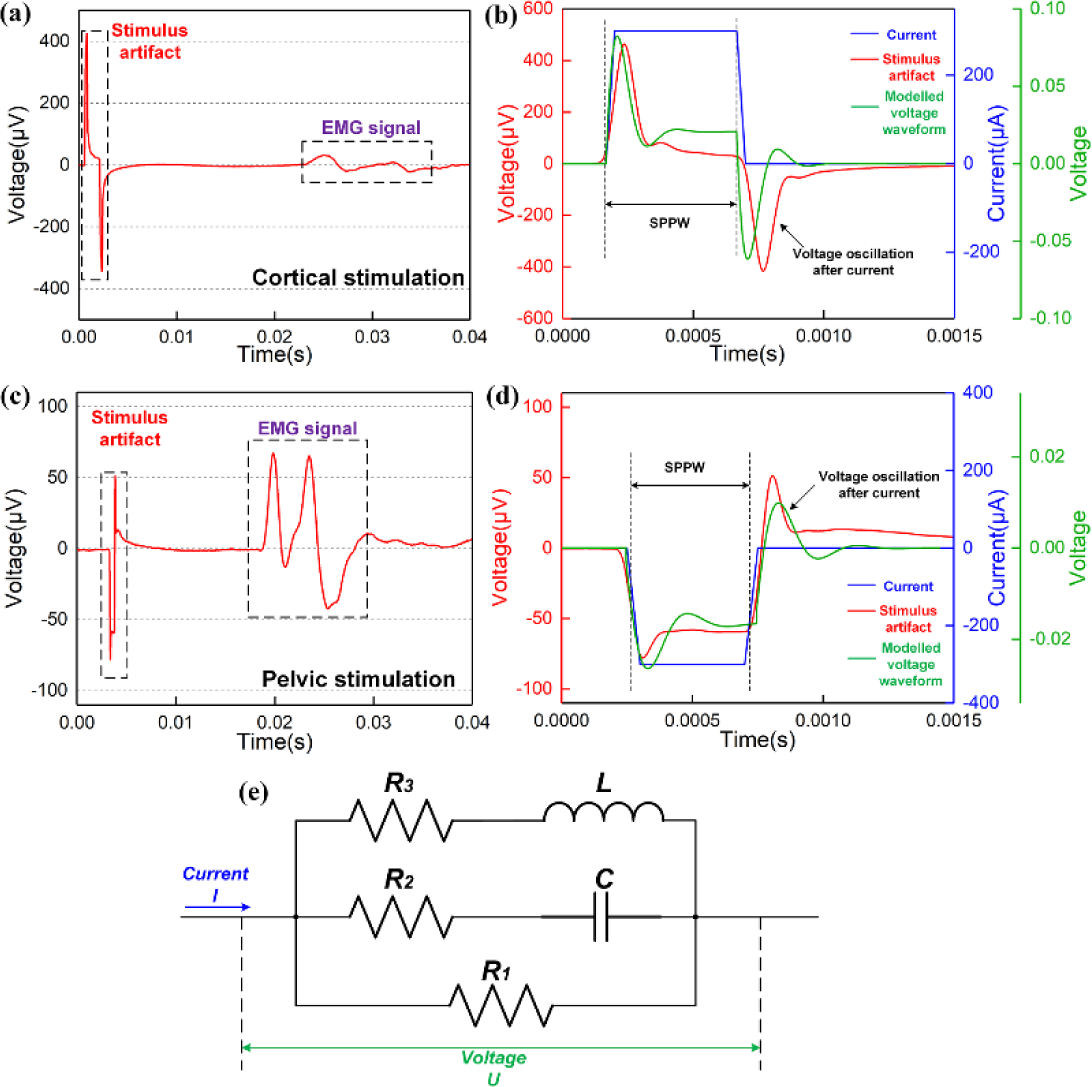
The stimulus artifacts in EMG recording from electrical stimulation of the cortical and pelvic nerve can be fitted by the voltage response of a parallel RLC circuit. (a) An EMG signal sample of the cortical stimulation; (b) The detailed analysis of the stimulus artifact: the recorded stimulus artifact is the red curve, the applied current is the blue curve and the voltage response of the parallel RLC circuit by modelling is the green curve; (c) An EMG signal sample of the pelvic nerve stimulation; (d) The detailed analysis of the stimulus artifact: the recorded stimulus artifact is the red curve, the applied current is the blue curve and the voltage response of the parallel RLC circuit by modelling is the green curve; (e) The circuit configuration of the parallel RLC circuit used for modelling.

A detailed analysis of the stimulus artifacts are shown in Figure 1(b) and (d), respectively. A monophasic square wave current pulse (blue curve) of a certain single phase pulse width (SPPW) was applied. As seen, the stimulus artifacts (red curve) and the current pulse share the same rising edge and falling edge. But there are two interesting points to be observed:

1. Within the current pulse, the stimulus artifact has a certain voltage oscillation.
2. After the current pulse, there is another voltage oscillation with an opposite polarity.

Previously, it was widely believed that there is no inductance in the neural circuit. The whole neural circuit should follow a general RC configuration. However, the electrical response of the RC circuit to a square wave current pulse will not show a voltage oscillation. This voltage oscillation is the feature of an RLC circuit. Therefore, we tried fitting the stimulus artifact with an RLC circuit. Here we used a parallel RLC circuit (Figure 1(e)) for modeling to fit the stimulus artifacts, shown as green curves in Figure 1(b) and (d), respectively. The voltage oscillations within and after the current pulses can be duplicated and the whole waveform can be generally fitted. The detailed modeling parameters in Figure1(b) and (d) of all electrical units can be found in **Table 1(2-(b)&(d))**.

**Table 1.** Modelling parameters

| No. | $R_1(\Omega)$ | $R_2(\Omega)$ | $R_3(\Omega)$ | $C(F)$ | $L(H)$ |
| --- | --- | --- | --- | --- | --- |
| 2-(a) | 2000 | 1350 | 500 | 10n | 0.1464 |
| 2-(b) | 3701 | 350 | 500 | 10n | 0.1464 |
| 2-(c) | 9000 | 1350 | 500 | 10n | 0.2326 |
| 2-(d) | 80000 | 300 | 1700 | 18n | 0.1086 |
| 2-(e) | 2656 | 1800 | 800 | 18n | 0.0813 |
| 3-(f) | 2656 | 1800 | 800 | 18n | 0.0813 |
| 8-(b) | 16579 | 100 | 3000 | 12n | 2.1109 |

From the waveform fitting in Figure 1(b) and (d), it is quite possible that the general equivalent neural circuit should follow a parallel RLC configuration. We can validate this from two aspects:

1. By changing the SPPW and the waveforms of the current pulses, the waveforms of the stimulus artifacts will change with a certain pattern, which can also be fitted by the electrical voltage response of the equivalent parallel RLC circuit. In other words, the stimulus artifacts generated by any kinds of current waveforms with any pulse widths shall be fitted.
2. The voltage response of the parallel RLC circuit is frequency dependent. In some cases, when the quality factor, which is the Q factor, is high enough, there will be a resonance effect, resulting in a maximum amplitude of the recording data. So in both the stimulus artifact the EMG recording, there should be a measurable resonance frequency of the neural system.

The results of the validation of the two points mentioned above are shown in Figure 2. Three kinds of commonly used square current waveforms are used in tests: positive monophasic, negative monophasic and positive-first biphasic waveform (indicated at the left upper corner of all (i) figures). The SPPW is changed within a certain range to clearly show the changing pattern of the stimulus artifacts (all (i) figures). Then the modeling results are shown in all (ii) figures. All the modeling parameters can be found in **Table 1(2-(a) to (f))**.

**Fig. 2.**
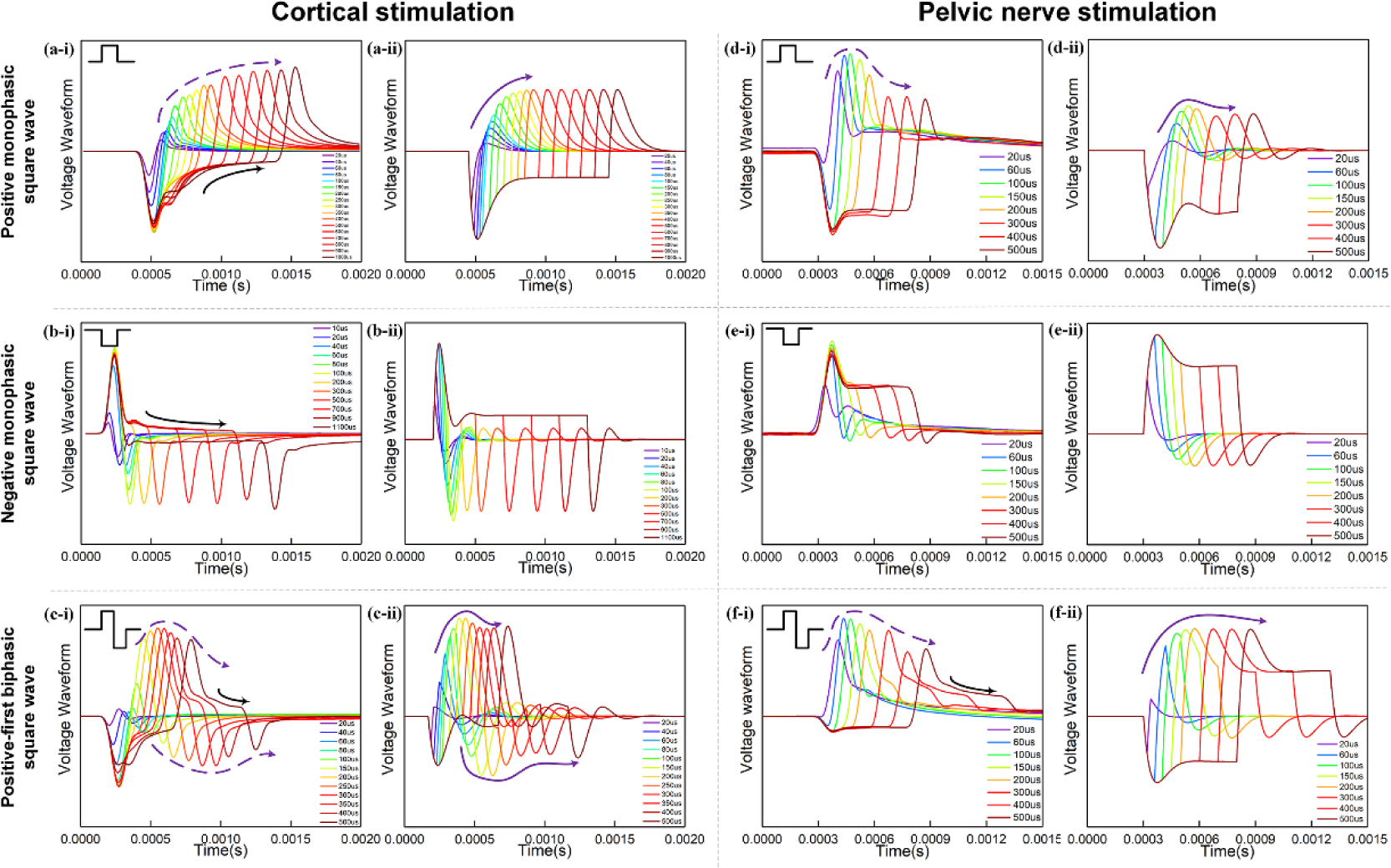
Experimental measurement and modelling (notation ‐i and ‐ii, respectively) of the stimulus artifact of cortical stimulation (a to c) and pelvic nerve stimulation (d to f) with different current waveforms and SPPW. (a-c) Cortical stimulation and modeming results: (a) positive monophasic square wave, (b) negative monophasic square wave and (c) positive-first biphasic square wave; (d-f) Pelvic nerve stimulation and modeming results: (d) positive monophasic square wave, (e) negative monophasic square wave and (f) positive-first biphasic square wave; (i) left figures refer to the measured data, (ii) right figures refer to the modelling results. The modelling results match well with the measurement data, validating the parallel RLC circuit used in this study.

Firstly, it can be clearly observed that the general changing pattern of the stimulus artifacts can be well fitted by the modeling results. Even the changing pattern of different waveforms are completely different, the changing pattern can always be duplicated by tuning the circuit parameters. Secondly, in Figure 2(c), (d) and (f), a distinct resonance effect can be found, indicated by the solid purple arrow in experimental data and dash purple arrow in modelling results. In cortical stimulation, the resonance effect happens between SPPW of 300 µs and 350 µs, referring to a resonance frequency between 1429 Hz and 1667 Hz. In pelvic nerve stimulation, the resonance effect happens between SPPW of 60 µs and 100 µs, referring to a resonance frequency between 5000 Hz and 8333 Hz.

An interesting phenomenon to be observed in Figure 2 is that, together with the RLC circuit response, some data shows an RC discharging pattern, indicated with the black arrows in Figure 2(a-i), (b-i), (c-i) and (f-i). This RC discharging effect can be recognized by comparing the experimental data with the modelling results. The voltage curve gradually drops at the position, whereas the voltage is predicted to keep constant in the modelling. This indicates that an additional capacitor should be added in the circuit to improve the modeming precision. The corresponding circuit with this additional capacitor and be found in the **supplementary S3**. And more repeated testing data showing the similar result can be found in the **supplementary S4**, validating the stability of the RLC neural circuit for analysis.

### B. Analysis of the EMG signals

The above-mentioned data confirms the existence of the inductance in the neural system. Then this inductance will also induce a resonance effect on the neural responses, which shall be observed in EMG signals. However, not all kinds of waveforms are effective to induce the nerve stimulation, only the data with successful stimulation is shown in Figure 3 and 4.

**Fig. 3.**
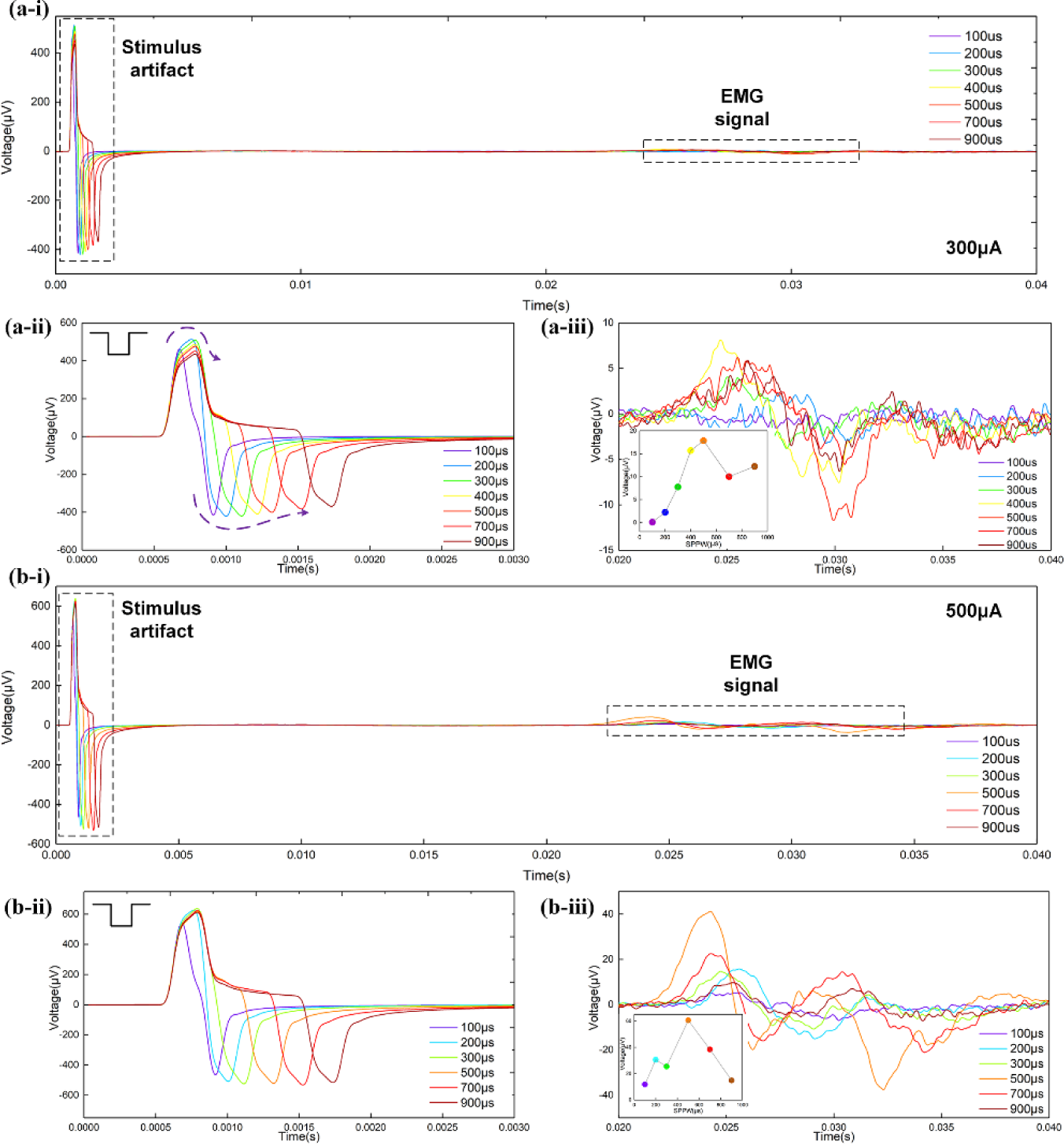
EMG recording of cortical stimulation by applying negative monophasic square current pulses of (a) 300 µA and (b) 500 µA: (i) The complete EMG data including the stimulus artifacts and EMG signal; (ii) detailed stimulus artifacts; (iii) detailed EMG signals, inset figure shows the voltage amplitude of EMG signals with SPPW.

The cortical stimulation results are shown in Figure 3. The negative monophasic square current pulses of 300 µA and 500 µA with SPPW from 100 µs to 900 µs were applied. Figure 3(a-i) and (b-i) shows the complete data. The detailed artifacts are shown in Figure 3(a-ii) and (b-ii), respectively. As seen, these stimulus artifacts share the same pattern as observed in Figure 2(b-i). The detailed EMG signals are shown in Figure 3(a-iii) and (b-iii). The inset figures show the amplitude of the EMG signal with different SPPWs. The resonance effect is quite distinct. The amplitude peaks at SPPW= 500 µs for both results of 300 µA and 500 µA, indicating that the resonance frequency of cortical is 1000 Hz.

The pelvic nerve stimulation results are shown in Figure 4. Three kinds of waveforms were applied with a constant current amplitude, 70 µA. The complete data is shown in Figure 4(a). The detailed stimulus artifacts can be found in Figure 2(d-f). The EMG amplitudes with SPPW for three waveforms in Figure 4(b) shows a clear resonance effect happened at SPPW=150 µs, which agrees with the resonance frequency of the stimulus artifact. The resonance effect is the strongest for the negative monophasic waveform: the amplitude of the EMG signal significantly drops when the SPPW is higher than 150 µs.

**Fig. 4.**
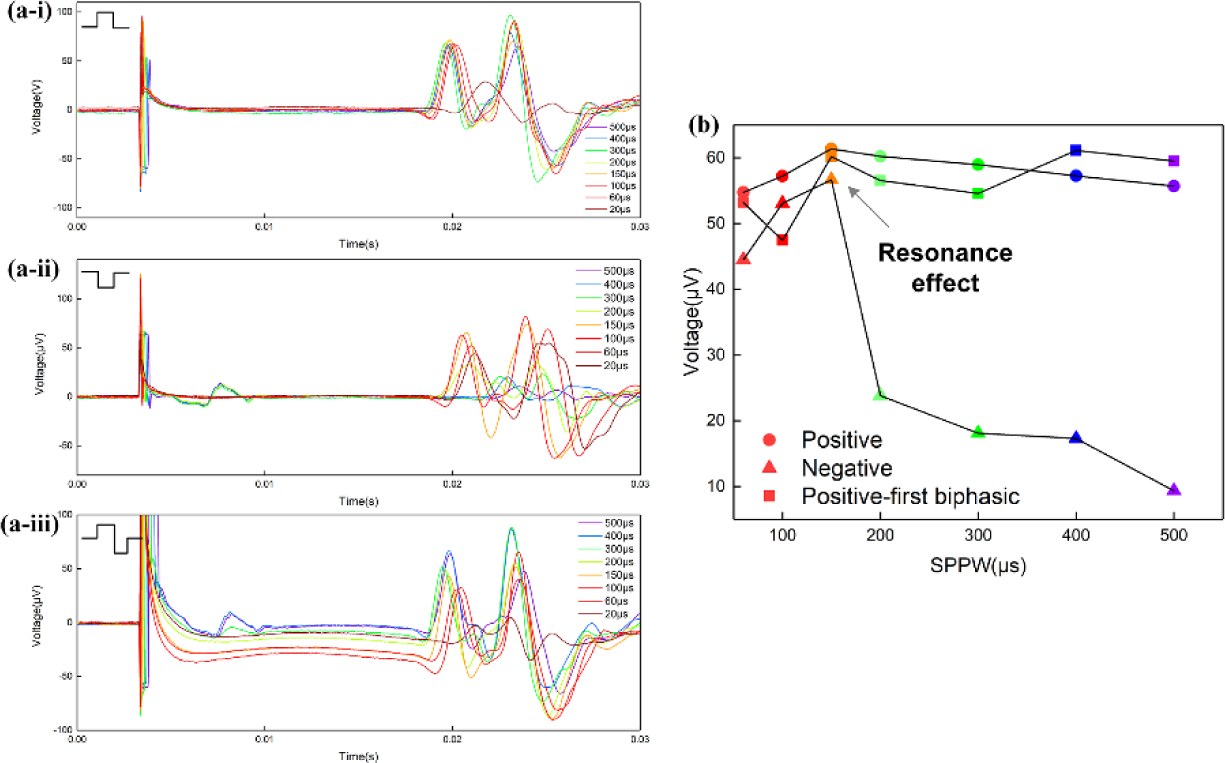
EMG recording of pelvic nerve stimulation by applying (a-i) positive monophasic, (a-ii) negative monophasic and (a-iii) positive-first biphasic square current pulses of 70 µA; (b) The voltage amplitude of EMG signals of three waveforms with SPPW.

Up to here, we have shown clear data to prove the existence of the inductance in neural tissue. People may

1. Why the resonance frequencies of the stimulus artifacts and EMG signal are not matched with each other. In pelvic nerve stimulation, the resonance frequencies of stimulus artifacts and EMG signal are all within the range from 5000 Hz to 8333 Hz. But in cortical stimulation, the resonance frequency of stimulus artifacts is between 1667 Hz and 1429 Hz (Figure 2(a-c)), which is quite different from the resonance frequency in EMG signals, which is 1000 Hz.
2. The exact circuit configuration of the real neural tissue should be very complicated, which normally be considered as a distributed parameter circuit. Why can we model it with a simple parallel RLC circuit?

These two questions will be answered in **Supplementary S5**.

### C. Derivation of the Strength-duration relationship

Another evidence to support the parallel RLC circuit configuration comes from a well-known empirical model, which called the strength-duration relationship. Now, this model can be directly derived and moreover, can be amended to a more precise format.

Previously, it is widely believed that charge is the factor to induce nerve stimulation. In such charge based theory, there is an empirical linear relationship between the threshold charge level and pulse duration, which is called Weiss’s strength-duration equation *(28)* for negative monophasic square current pulses. This equation describes the threshold charge as a function of pulse width as follows:

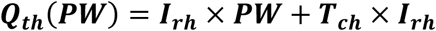

where ***I_rh_*** is the rheobase current, ***T_ch_*** is the chronaxie, and ***PW*** is the pulse width. The rheobase current is defined as the threshold current for infinitely long pulses. The chronaxie is defined as the pulse duration required for excitation when the current amplitude is equal to twice the rheobase current. And Lapicque reiterated Weiss’s equation for the strength–duration relationship *(29)*, but in terms of the threshold current, and introduced the rheobase current and chronaxie as the constants:

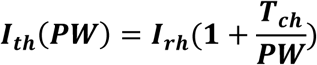

Apparently, these two equations are just mathematical descriptions without explaining how ***I_rh_*** happen and why the curve follows a specific trend. As follows is the derivation of this relationship with the physical definition of ***I_rh_***.

Figure 5(a) shows a typical voltage waveform by applying negative monophasic square current pulses with different SPPW. For the voltage waveform of each SPPW, the peak voltage is denoted as ***V_p_***, which is a function of the current, ***I***, and the pulse width, ***SPPW***, written as ***V_P_***(***I***, ***SPPW***). Here we need to set a hypothesis that the nerve stimulation happens when the voltage exceeds a threshold voltage, which is denoted as ***V_Threshold_***. This ***V_Threshold_*** is of a negative value since the ion channel can be activated by a negative voltage. Thus the criteria to induce the nerve excitation is ***V_P_***(***I***, ***SPPW***) ≥ ***V_Threshold_***. Then both the threshold current, ***I_th_***, and the threshold charge, ***Q_th_*** = ***I_th_*** × ***SPPW***, are defined as the current and charge required to make the ***V_P_*** reaches ***V_Threshold_***.

**Fig. 5.**
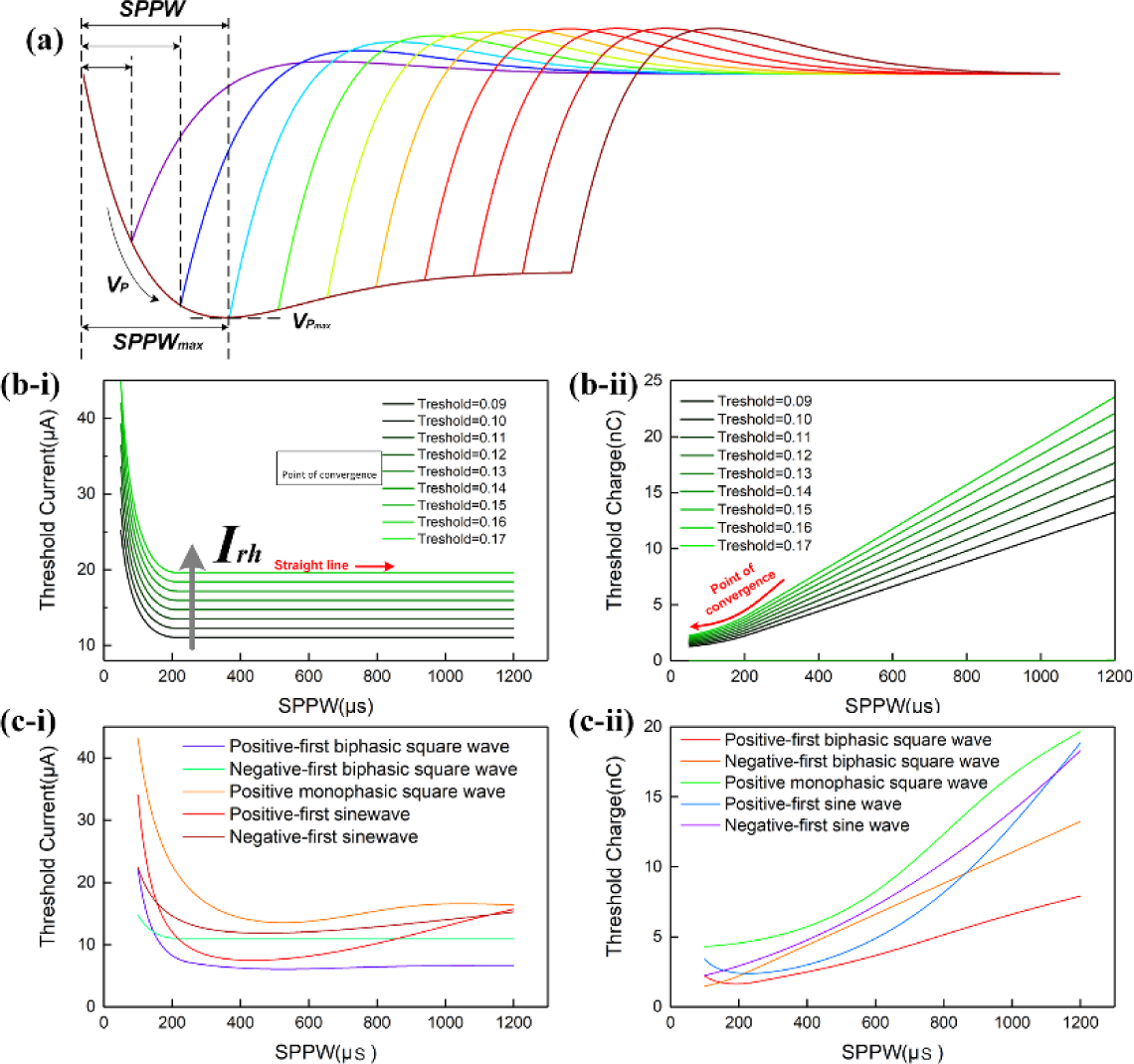
Derivation of the Strength–duration relationship. (a) Illustrative voltage waveforms generated by negative monophasic square waveform current; (b-i) The threshold current amplitude (***I_th_***) decreases as the ***SPPW*** increases in a nonlinear fashion; (b-ii) The relationship between threshold charge (***Q_th_***) and ***SPPW*** is linear; (c-i) The relationship between the threshold current amplitude (***I_th_***) and the ***SPPW*** for different current waveforms; (c-ii) The relationship between threshold charge (***Q_th_***) and ***SPPW*** for different current waveforms.

Then the critical condition is:

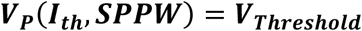

***I_th_*** and ***Q_th_*** can be written as functions of ***SPPW*** and ***V_Threshold_***:

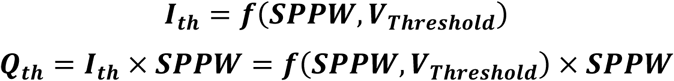

Since ***V_p_***(***I_th_***, ***SPPW***) cannot be expressed analytically, only numerical solutions of ***I_th_*** and ***Q_th_***, which are calculated with a set of modeling parameter, are provided in Figure 5(b) and Figure 5(c). In Figure 5(b), all curves decrease to a constant value, ***I_rh_***. This is because the ***V_P_*** will saturate at a maximum value, 
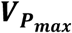
, when 
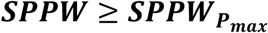
, as shown in Figure 5(a).

Meanwhile,

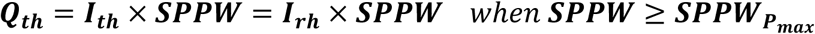

Since ***I_rh_*** is a constant, ***Q_th_*** increases linearly with ***SPPW***, when 
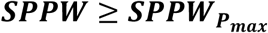
, as shown in Figure 5(c).

The physical meaning of ***I_rh_*** is the threshold current when 
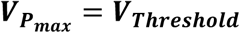
. Meanwhile, the nonlinear curve of ***I_th_*** versus ***SPPW***, the existence of ***I_rh_*** and linear curve of ***Q_th_*** versus ***SPPW***, can be directly obtained without any additional hypotheses. The exact analytical equation for this relationship is not available. The curves in Figure 5(b) and Figure 5(c) are the numerical solutions of strength-duration relationship. It corrects the relationship in two aspects:

1. Rather than infinitely approaching to the ***I_rh_*** as the case in Weiss’s strength–duration equation, the threshold current curve will be equal to the ***I_rh_*** when 
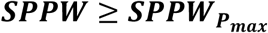
, which is a straight line indicated by the black arrow in Figure 5(b).
2. Rather than being a completely straight line, the threshold charge curve is linear only when 
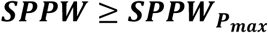
. When the ***SPPW*** is approaching zero, the slope of the threshold charge curve will also approach zero, meaning that the threshold charge will converge in a constant value at low ***SPPW***, indicated as the point of convergence in Figure 5(c).

These two major special differences with the Weiss’s equation have already be confirmed by previous research *(30,31)* and now can be well explained with the parallel RLC circuit model.

Moreover, it also explains why such a relationship is only valid to the negative monophasic square current waveform. Because the voltage waveforms differ with the current waveforms, inducing a more complicated trend without a stable ***I_rh_***, which was observed in other researches *(32)*. In Figure 5 (d) and (e), representative strength-duration curves of other waveforms including different types of square waves and sine waves are shown. As seen, these curves don’t share the same curve pattern as that of the negative monophasic square current pulse. For example, the threshold current of sinewave current increases at high SPPW range, this phenomenon has been observed in previous research with triangle current waveform *(32)*. But as explained before, the analytical solution for this relationship is not available, we can only numerically calculate the curve, which highly depends on the exact circuit parameter. Therefore, except the case of the negative monophasic square wave, the exact curve shape of other waveforms may vary with different circuit parameter. This explains why this relationship valid for negative monophasic square current pulses.

## Theoretical explanation of the physical entities for the inductance

### A. The inductance generated by the coil structure of myelin sheaths

The biological structure of a segment of the myelinated axon is shown in Figure 6(a). The proposed distributed-parameter circuit model is as shown in Figure 6(b), which is equivalent to a segment of a single axon shown in Figure 6(a). This axon segment has two parts with different structures: part with myelin sheath and node of Ranvier without myelin sheath. For the Ranvier node, the cell membrane can be modelled as capacitors, ***C*_1_**, connecting the inside terminal and outside terminal of the cell membrane. The intracellular and extracellular fluid can be modelled as resistors. For the part covered by myelin sheath, an extra inductors, ***L***, is connected between the cell membrane, ***C*_2_**, and extracellular fluid. Then the whole axon can be modelled as a circuit cascade as shown in Figure 6(c). Multiple axons can be modelled as a cascade network as shown in Figure 6(d). The outside terminals of each axon are connected with resistors. Meanwhile, a key feature need to be added in this circuit, which is the mutual inductance between the adjacent myelin sheaths. In term of physics, this mutual inductance can be either positive or negative, depending on their mutual spiraling orientations. In this circuit model, a definite conclusion will be drawn that this mutual inductance can only be positive (will be explained in the next section), meaning that the spiraling orientations of adjacent myelin sheaths have to be opposite. We noticed that this phenomenon has been reported by previous experimental observations *(33-35)* but cannot be explained by any previous theories and models.

**Fig. 6.**
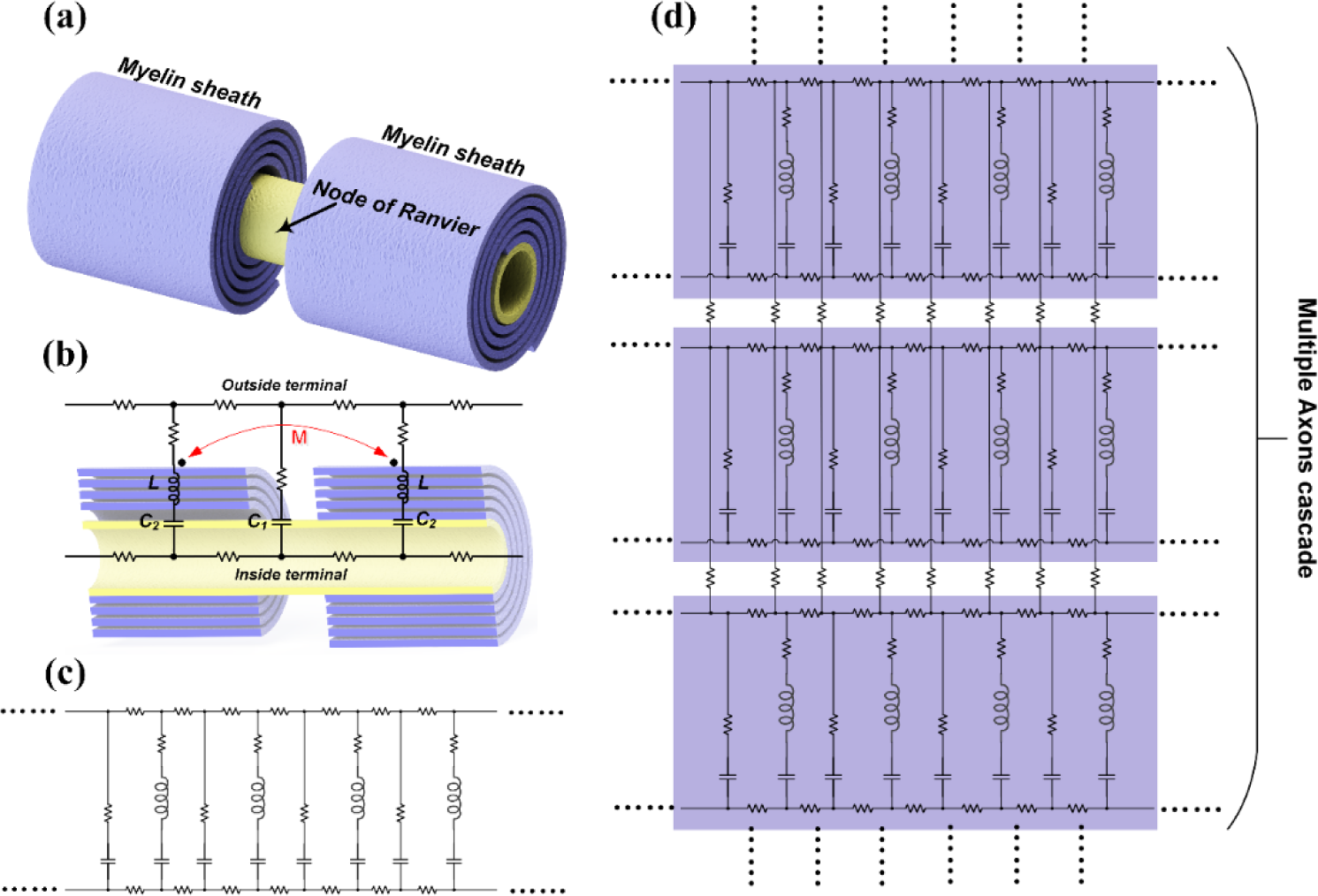
The equivalent neural circuit of axons by considering the myelin sheaths as inductors. (a) 3D illustration of the biological structure of a myelinated axon segment; (b) the cross-sectional view of the axon segment with the equivalent circuit model; (c) the equivalent circuit cascade of a myelinated axon; (d) the equivalent cascaded circuit network of multiple axons.

#### 1. How the distributed-parameter circuit compatible with the lumped-parameter circuit

A single axon can be modeled as a cascade of basic units, indicated within the red dash box in Figure 7(a)). The configuration of this basic unit is a parallel RLC circuit with an extra capacitor (***C*_2_**) connected in series with the inductor. Here the physical meaning of ***C*_1_** is the capacitance of the Ranvier node and ***C*_2_** is the capacitance of the cell membrane covered by myelin sheath. Considering that the length of the segment covered by myelin sheath is much longer than the Ranvier node, here ***C*_2_** ≫ ***C*_1_**. Apparently, this cascade can be simplified as a circuit which has the same configuration as its basic unit, as shown in Figure 7(c).

**Fig. 7.**
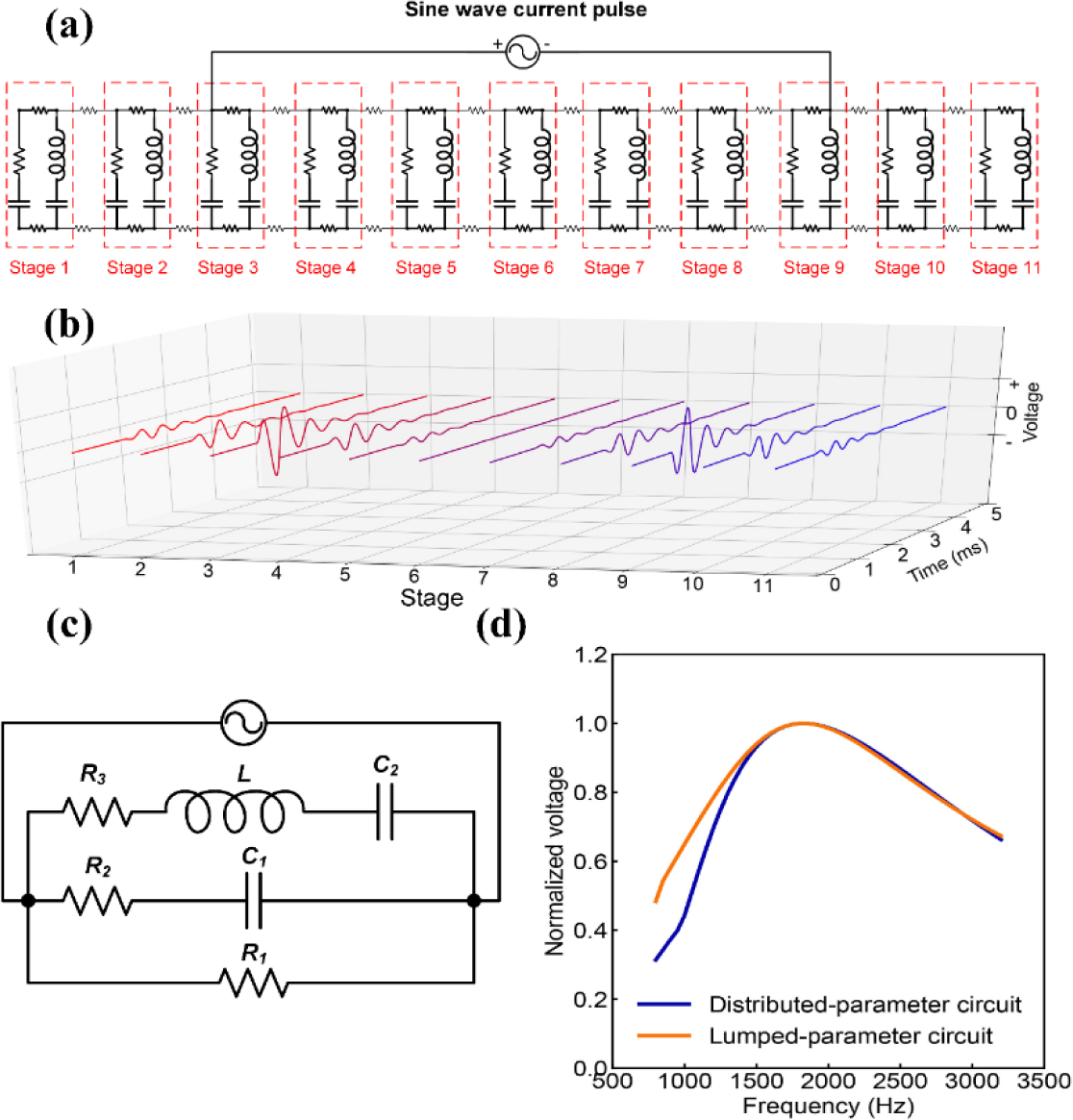
Circuit analysis showing the lumped-parameter circuit can used as simplification of the distributed-parameter neural circuit. (a) A circuit cascade with 11 stages for modeling; (b) The voltage waveforms on ***C*_1_** of each stage, showing a gradual change of the amplitude the polarity reversing; (c) The simplified lumped-parameter circuit proposed in previous experimental analysis; (d) The frequency response curve of the voltage amplitude on ***C*_1_** of the stage 4 in (a) and (c).

This simplified circuit is exactly the revised lumped-parameter circuit proposed in Figure S3. In the initial version of the lumped-parameter circuit shown in Figure 1, the extra capacitor ***C*_2_** is not included. However, in Figure 2, we have already shown that an extra capacitor should be added for a better curve fitting. Now based on the distributed-parameter circuit, which is derived from the biological structure of the axon, we know this ***C*_2_** really exists and its physical meaning is the cell membrane covered by the myelin sheath.

This circuit cascade can be directly used for the study of the nerve stimulation. An equivalent circuit cascade with 11 stages is shown in Figure 7(a). A current source is connected to the stage 3 and 9 (Figure 7(a)), which is the similar situation as an electrode pair implanted on a nerve branch. When a sinewave current pulse is applied, the voltage waveforms on the capacitors, ***C*_1_**, of each stage are shown in Figure 7 (b). As can be seen, the voltage waveforms will gradually change from a positive-first phase to a negative-first phase. It means the segment close to the positive electrode actually has a different voltage waveform with the segment close to the negative electrode. As a simple prediction, with a same stimulating current waveform, the stimulation to the upstream and downstream will be different. With correct circuit parameters, this difference shall be well explained by the polarity reversing of the voltage waveform.

Then a typical case demonstration in Figure 7(d) shows how the lumped-parameter circuit (Figure 7(c)) generates the same electric characteristics as the distributed-parameter circuit (Figure 7(a)), validating the circuit simplification proposed in this study. With the same configuration as shown in Figure 8(a), by varying the frequency of the sinewave current pulse while keeping the amplitude the same, the frequency response of the voltage amplitude on ***C*_1_** of stage 4 is shown as the blue curve in Figure 8(d), which is a typical frequency response curve of a parallel RLC circuit. Then by tuning the circuit parameters in Figure 7(c), a similar frequency response of the voltage amplitude on ***C*_1_** in the lumped-parameter circuit (Figure 7(c)) is shown as the orange curve in Figure 7(d), which is almost the same as the blue curve.

**Fig. 8.**
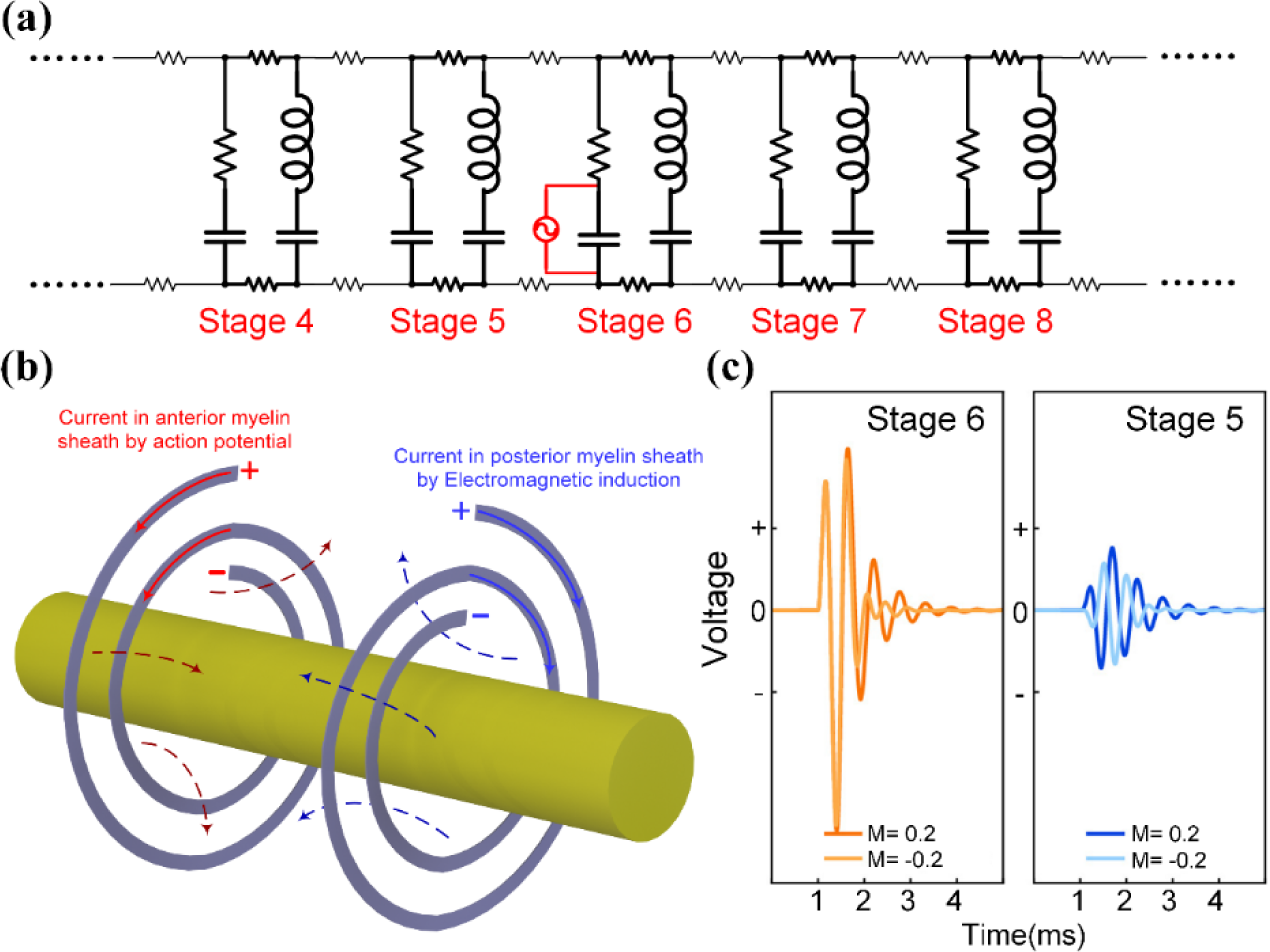
Circuit analysis showing the opposite wrapping orientations of adjacent myelin sheaths can enhance the signal propagation speed. (a) A circuit cascade with 11 stages for modeling, the current source for applying a sine wave current pulse is connected on ***C*_1_** of stage 6 to mimic the activation of an action potential; (b) Illustrative plotting of the mutual inductance between adjacent myelin sheaths; (c) Comparison of the voltage waveforms on ***C*_1_** of stage 5 and 6 with positive and negative mutual inductance.

In summary, if we only care about the voltage waveforms on the Ranvier node, which determines the result in electrical nerve stimulation, a lumped-parameter circuit in Figure 7(c) can simplify the analysis. However, an in-depth study of the distributed-parameter circuit can help understand more about the nervous system, which will be explained in the following sections.

#### 2. How the myelin sheath enhances the propagation speed of neural signals?

In conventional neural models, the myelin sheath is always modeled as a pure resistor to increase the propagation speed of neural signals by decreasing <u>capacitance</u> and increasing <u>electrical</u> <u>resistance</u> across the cell membrane. However, in our new circuit model by considering the myelin sheath as an inductor, this propagation speed enhancement will have a new explanation. Generally speaking, this propagation speed enhancement is induced by two mechanisms: mutual inductance between two adjacent myelin sheaths and frequency modulated signal decay. To demonstration these two mechanisms, a circuit which is similar to one in Figure 7(a) is proposed in Figure 8(a), with a different configuration of the current source. The current source is connected in parallel with the ***C*_1_** of stage 6. A sinewave current pulse is applied to mimic the generation of action potential.

##### a. Mutual inductance

Since the node of Ranvier is very short, the mutual inductance between the two adjacent myelin sheaths will not be negligible. An illustrative plotting for this mutual inductance is shown in Figure 8(b). When an action potential is activated on the node of Ranvier before the anterior myelin sheath, this action potential is also coupled onto this myelin sheath. Assuming at an instant, the potential cross the myelin sheath is positive outside and negative inside, then a current will flow along the spiraling orientation from outside to inside, generating a magnetic field. This magnetic field will generate an inductive current on the posterior myelin sheath. Due to the opposite spiraling orientations of adjacent myelin sheaths, the voltage induced by the inductive current on the posterior myelin sheath will share the same polarity, which is also positive outside and negative inside. This voltage will be further coupled onto the node of Ranvier between these two myelin sheaths. Surely, even without this mutual inductance, the action potential on the anterior node of Ranvier still can generate some voltage onto the posterior node of Ranvier. However, this mutual inductance can increase the voltage amplitude, reduce the signal decay and finally enhance the propagation speed of neural signal.

The effect of the mutual inductance upon the propagation of the action potential is shown in Figure 8(c). A sine wave current pulse is applied on ***C*_1_** of stage 6 to mimic the generation of an action potential. The voltage waveforms on ***C*_1_** of stage 6 and stage 5 are compared when the mutual inductance is positive (opposite spiraling orientations) and negative (same spiraling orientations). This mutual inductance only has a minor effect on the voltage of stage 6, but affects the voltage of stage 5 a lot. The voltage of positive mutual inductance is higher than that of negative mutual inductance. Meanwhile, the positive mutual inductance can guarantee a same voltage polarity between the anterior and posterior stage, which can make the activation of the action potential on the posterior stage earlier. But the negative mutual inductance may induce a polarity reversing. As shown in Figure 8(c), the voltage on stage 6 has a positive-first polarity while the voltage for negative mutual inductance on stage 5 has a negative-first polarity, which will delay the activation of action potential.

##### b. Frequency modulated signal decay

Another effect induced by myelin sheaths upon the neural signal propagation is the frequency modulated signal decay. Considering the same situation as shown in Figure 9(a), an action potential is activated on the stage 6, the voltage waveforms on ***C*_1_** of each stage are shown in Figure 9(a). The voltage amplitude reaches maximum on stage 6 and gradually decay for the stages farther from stage 6. In this circuit, the decay constant ***λ*** is defined as the voltage ratio between the anterior and posterior stages:

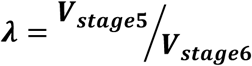

This decay constant ***λ*** is determined by the frequency of the input signal. A detailed circuit analysis can be found in **Supplement S6**. A simple conclusion drawn from the circuit analysis is that the decay constant ***λ*** reaches maximum at the resonance frequency of each stage, meaning a minimum signal decay. This conclusion is also confirmed by modeling as shown in Figure 9(b)-(d). The voltage amplitudes of each stage with different input frequency is shown in Figure 9(b). For the curve of 2 kHz, which is the resonance frequency set for each stage, the response voltage has the maximum amplitude and minimum decay. In Figure 9(c), the voltage-frequency curves of stage 5 and stage 6 are shown. These two curves all exhibit a parallel RLC circuit characteristic, which is similar to the result in Figure 7(d). Then calculate the decay constant ***λ*** in Figure 9(d). As can been seen, ***λ*** reaches maximum at the frequency very close to resonance frequency set for modelling, which is 2 kHz. But there is still a slight frequency mismatch. This is because the resonance frequency set for modelling is calculated by the equation:

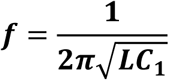

However, the actual resonance frequency is also affected by factors such as ***C*_2_**, other resistors and mutual inductance. So the frequency with maximum ***λ*** has a certain shift with 2 kHz. In this case, the actual resonance frequency is 1500 Hz. It means the action potential can propagate with a minimum decay when its major frequency is at 1500 Hz. Or in other words, the growth of the myelin will fit the major frequency of the action potential to realize a minimum signal decay.

**Fig. 9.**
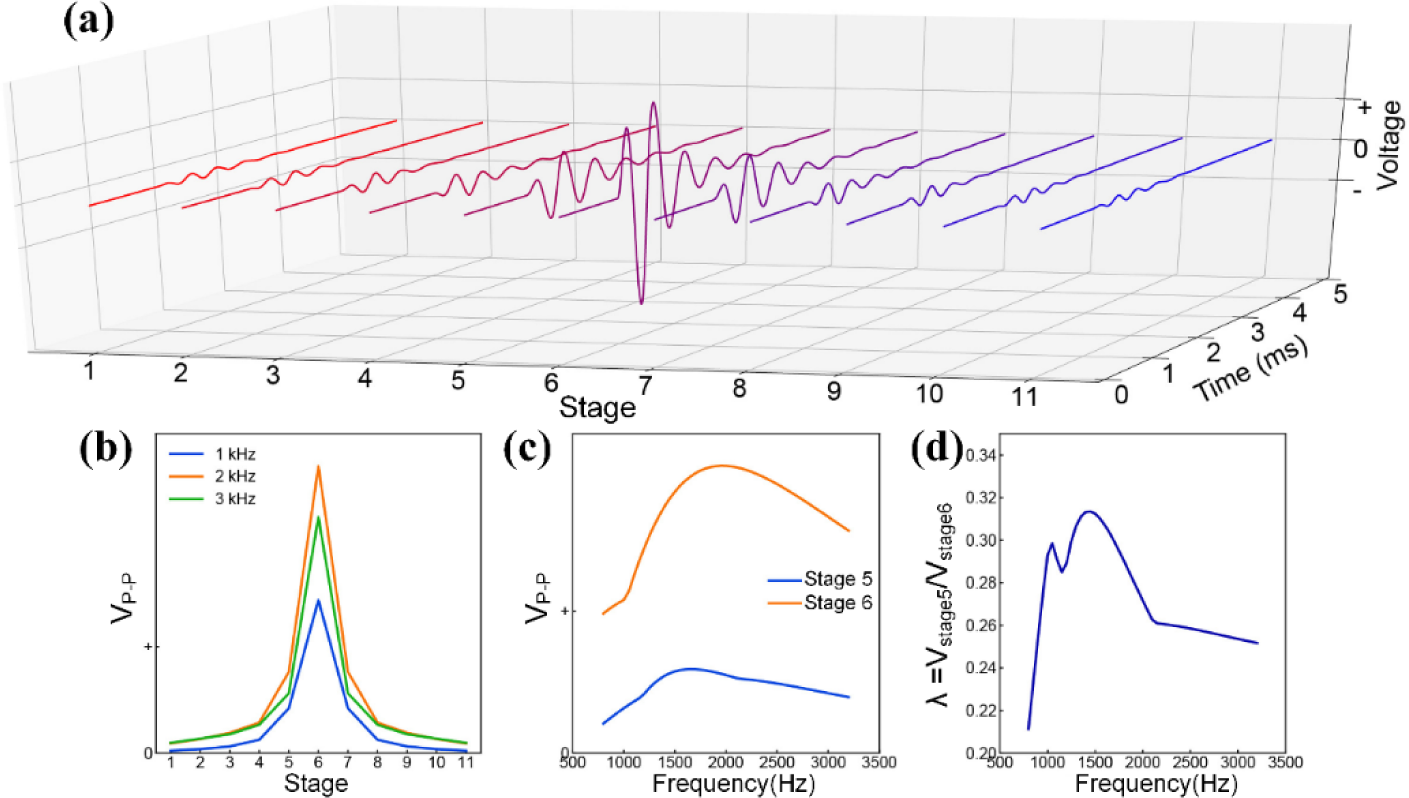
Circuit analysis showing myelin sheaths can minimize the spatial decay of the neural signal and enhance the propagation speed. (a) Voltage waveforms on ***C*_1_** of each stage when an action potential is activated on stage 6, the circuit for modeling is shown in Figure 8(a); (b) The amplitude of the voltage waveform on each stage with different frequency of the applied sine wave current pulse applied on stage 6; (c) The voltage-frequency curve on ***C*_1_** of stage 6 and stage 5; (d) the curve of decay constant 
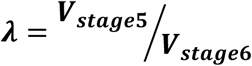
 with respect to frequency

In summary, the myelin sheath can enhance the propagation of neural signal by two mechanisms. The opposite spiraling orientations of adjacent myelin sheaths can induce a positive mutual inductance to increase the coupling between the anterior and posterior nodes of Ranvier, enhancing the propagation speed of action potential. The myelin will grow to a certain length and thickness to make the resonance frequency of the axon the same as the major frequency of the action potential, minimizing the signal decay along the axon to enhance the propagation speed of action potential. Surely, as a direct prediction, there should be a magnetic field accompany with the electric signal of the action potential. This magnetic field with action potential has been measured for more than half century *(39-52)*.

## Effect of myelin on magnetic nerve stimulations

Since the myelin sheath is an inductor in our theory, an external applied magnetic field can also generate inductive current within the myelin sheath and further generate voltage on the node of Ranvier to activate action potential. An illustrative drawing of this electromagnetic induction is shown in Figure 10(a). When a magnetic field is applied along the longitudinal direction of the axon, the inductive current on the anterior and posterior myelin sheaths will flow in opposite directions because of their opposite spiraling orientations. So the inductive potential on these two myelin sheaths will have opposite polarities. Since the outside terminals of these two myelin sheaths are connected, these two inductive potential will cancel with each other and finally, only their potential difference can be coupled onto the node of Ranvier in between. This phenomenon can be modeled by the circuit as shown in Figure 10(b). The inductive potential can be modeled as a voltage source connected in series with the inductor. Here only stage 5 and 6 are used for a simple demonstration. The voltage source of these two stages will be applied with sine wave current pulse of opposite polarities. The amplitudes of these two voltage sources are ***V*_1_** and ***V*_2_** for stage 5 and 6, respectively.

**Fig. 10.**
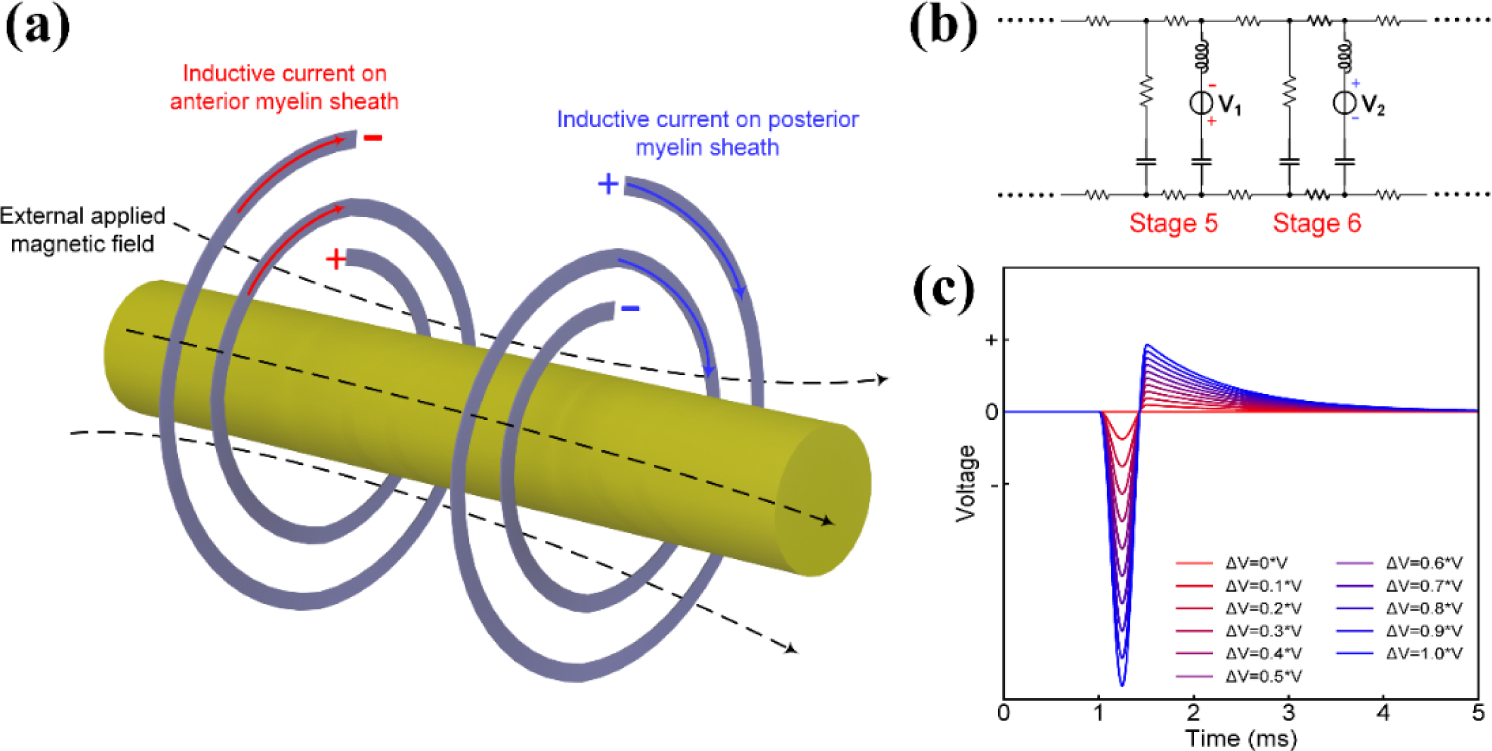
Circuit analysis showing the spatial gradient of the magnetic field determines the result of the magnetic nerve stimulation. (a) Illustrative drawing of the inductive current on adjacent myelin sheaths by external applied magnetic field; (b) Equivalent circuit with voltage sources connected to adjacent stages; (c) The voltage on ***C*_1_** of stage 6, representing the voltage on the node of Ranvier between two myelin sheath, by increasing the voltage difference between two stages.

The difference between ***V*_1_** and ***V*_2_** is *Δ**V***:

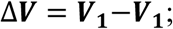

The voltage on ***C*_1_** of stage 6, which is the node of Ranvier between two myelin sheaths of stage 5 and 6, by increasing *Δ**V*** is shown in Figure 10(c). As can be seen, the voltage on ***C*_1_** is proportional to *Δ**V***. When there is no potential difference, the voltage on ***C*_1_** is also zero.

Since the inductive potential on the myelin sheath is proportional to the amplitude of the magnetic field component in the longitudinal direction of the axon, the voltage difference between two adjacent myelin sheaths is proportional to the amplitude difference of this magnetic field, which is the spatial gradient along the axon. It means, it is the spatial gradient, not the amplitude, of the magnetic field actually taking effect for magnetic nerve stimulation. This phenomenon is validated by previous study *(53,54)* and now can be explained by our theory. In normal condition when a coil is positioned near a straight axon, the gradient of the magnetic field along the axon is very low. It explains why the current required for magnetic field stimulation is so high *(53)*.

A more detailed analysis for nerves with different bending angles in magnetic stimulations is shown in Figure 11. Here only consider the situation when the magnetic field is along the axon. When the axon is straight (Figure 11(a)), the inductive potentials of the two near location are denoted as ***V*_1_** and ***V*_2_**. Since the magnetic field is along the axon, these two potential are all proportional to the amplitude of the magnetic field. Their difference is proportional to the gradient of the magnetic field:

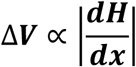

**Fig. 11.**
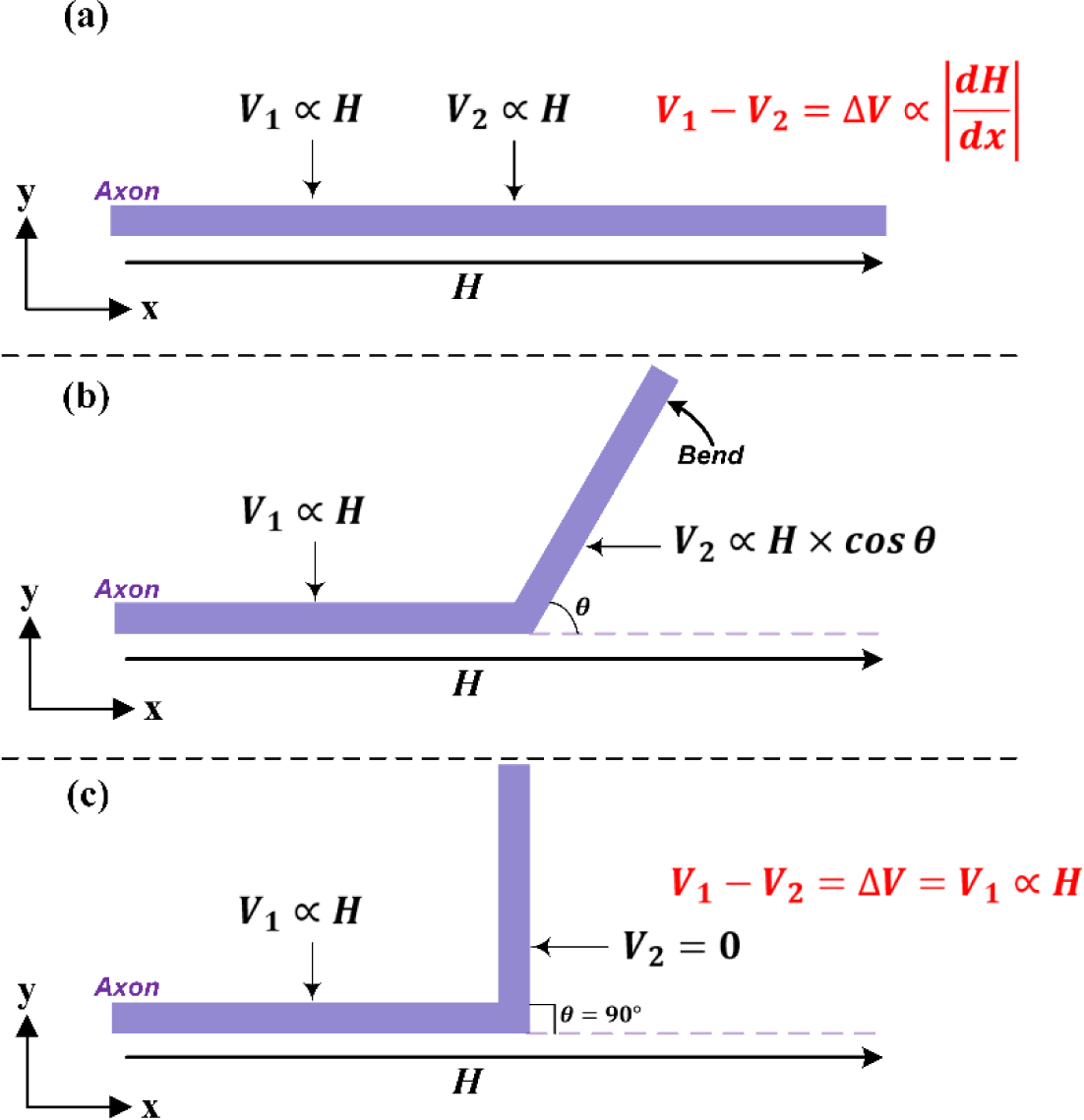
A detailed analysis of magnetic nerve stimulation with different axon bending angles. (a) The axon is straight; (b) The axon is bent with angle ***θ***; (c) The bending angle ***θ*** = **90**°.

The direction along the axon is set as ***x*** axis.

When part of the axon is bent with an angle ***θ*** as shown in Figure 11(b), the inductive potential ***V*_2_** will change as:

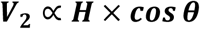

Generally, ***V*_2_** will decrease since ***cos θ*** ≤ **1**, and *Δ**V***, the difference between ***V*_1_** and ***V*_2_**, will increase, lowering the threshold current required for magnetic nerve stimulation. When the bending angle ***θ*** = **90**° as shown in Figure 11(c), ***V*_2_** = **0**. Then the voltage difference *Δ**V*** = ***V*_1_**, which is the theoretically maximum value can be achieved. In this situation, the current required for magnetic stimulation will be the minimum. The analysis results for these three situations in Figure 11 are exactly the experimental observations in previous study *(54)*. Now all these phenomena can be theoretically explained in our theory.

Moreover, another case demonstration in Figure 12 shows how our theory predicts an exact experiment observation in previous study *(54)*. It was observed that there will be two stimulation points when a straight nerve is placed above the coil with a figure of eight shape. Figure 12(a) shows the experiment configuration. A straight nerve along the x axis is placed above the coil and the current directions in two circles are anticlockwise and clockwise, respectively. Two yellow points refer to the two stimulation points. The magnetic field gradient along x axis, 
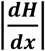
, in x-z plane when y deviates a bit from zero point, is shown in Figure 12(b). As can be seen, there will be two peaks, corresponding to the two stimulation points observed in previous study. It needs to be emphasized that the magnetic field along x axis is always zero when y=0, because of the perfect symmetric coil shape used in our modelling. The magnetic field along x axis generated by two circles will completed balance with each other. However, this situation will never happen in real experiment since the coils shape will never be perfectly symmetric and the nerve has a certain volume.

**Fig. 12.**
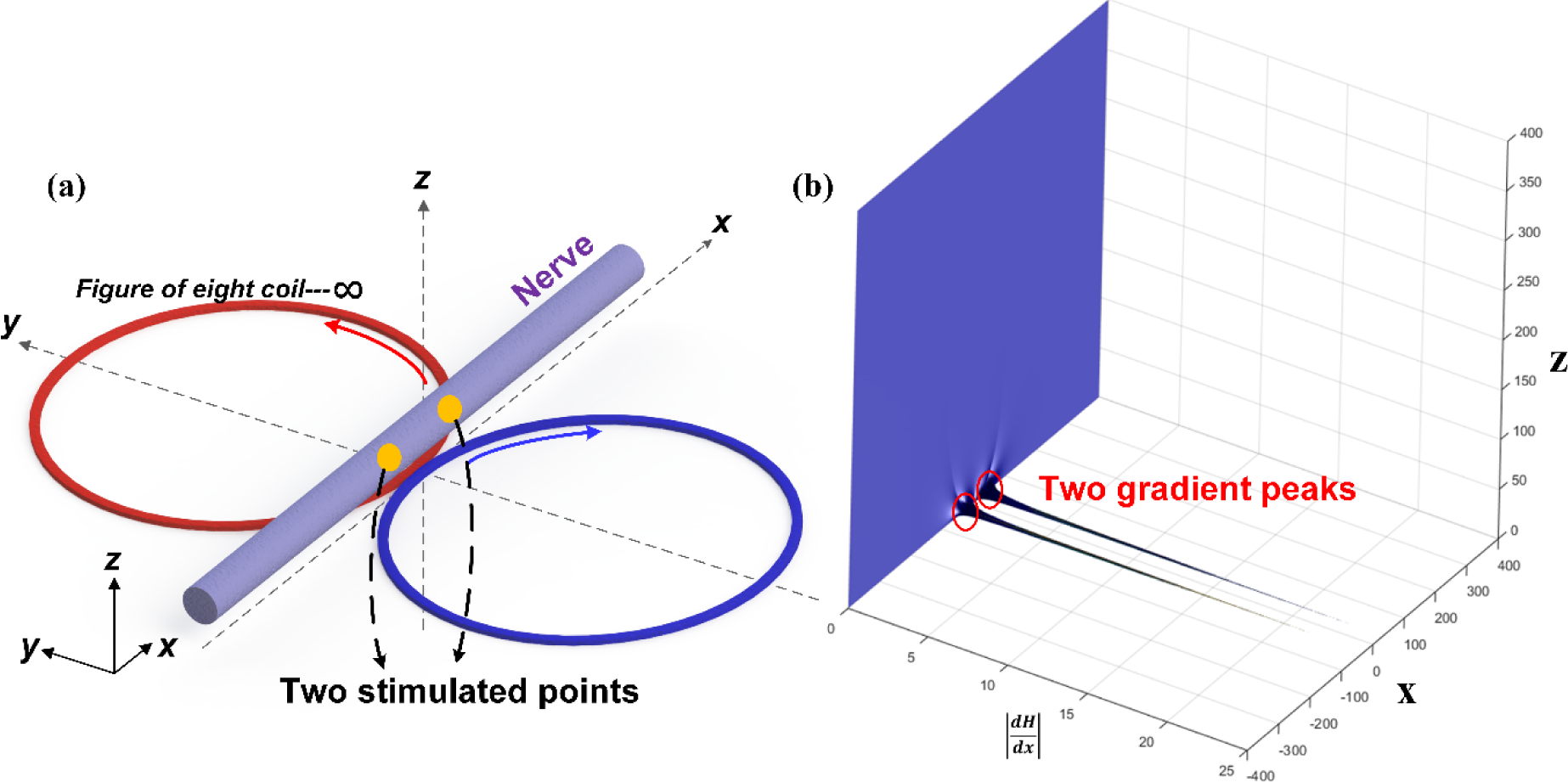
Modelling of magnetic field nerve stimulation by a figure of eight coil. (a) 3D illustrative drawing showing that a straight nerve along x axis is placed above a figure of eight coil, the current directions in two circles are anticlockwise and clockwise, respectively; (b) The modelling result of 
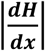
, the gradient of the magnetic field component along x axis, in x-z plane, showing two peaks corresponding to the two stimulation points observed in previous study *(25)*.

Since the coil structure is the main factor to enable the magnetic nerve stimulations, as a direct prediction, unmyelinated nerves and other excitable membranes such as muscle fibers will be very difficult to be stimulated by magnetic field. It is reported that C-fibers and cardiac nerve (unmyelinated nerve fiber) are hard to be stimulated by magnetostimulation *(55,56)*, which can be an evidence to support this prediction.

Based on the above explanation, it is clearly shown that the coil structure of myelin sheath can generate an inductive reactance to realize an energy conversion between electric field and magnetic field. In perspective of circuit, myelin sheath should be considered as a real inductor, which can generate magnetic field from electric current and sense magnetic field by generating inductive current. However, based on the equation of the resonance frequency:

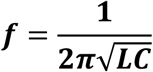

This inductance will be a huge value, can be as high as several Henry (H), when we consider the cell membrane capacitance as a reasonable value, which is of nF level, and the measured resonance frequency ***f*** is at kHz level. This huge inductor has been measured in the original paper of H-H model *(2)* and Cole’s early stage results *(3)*. Apparently, the myelin sheath, which is a microscale object, cannot provide such as a huge inductance. Thus, there should be another mechanism to provide this huge inductance. Actually, this huge inductor is not a real one, but an equivalent one. It is induced by the piezoelectric effect of the plasma membrane. This piezoelectric effect is illustrated in Figure 13.

**Fig. 13.**
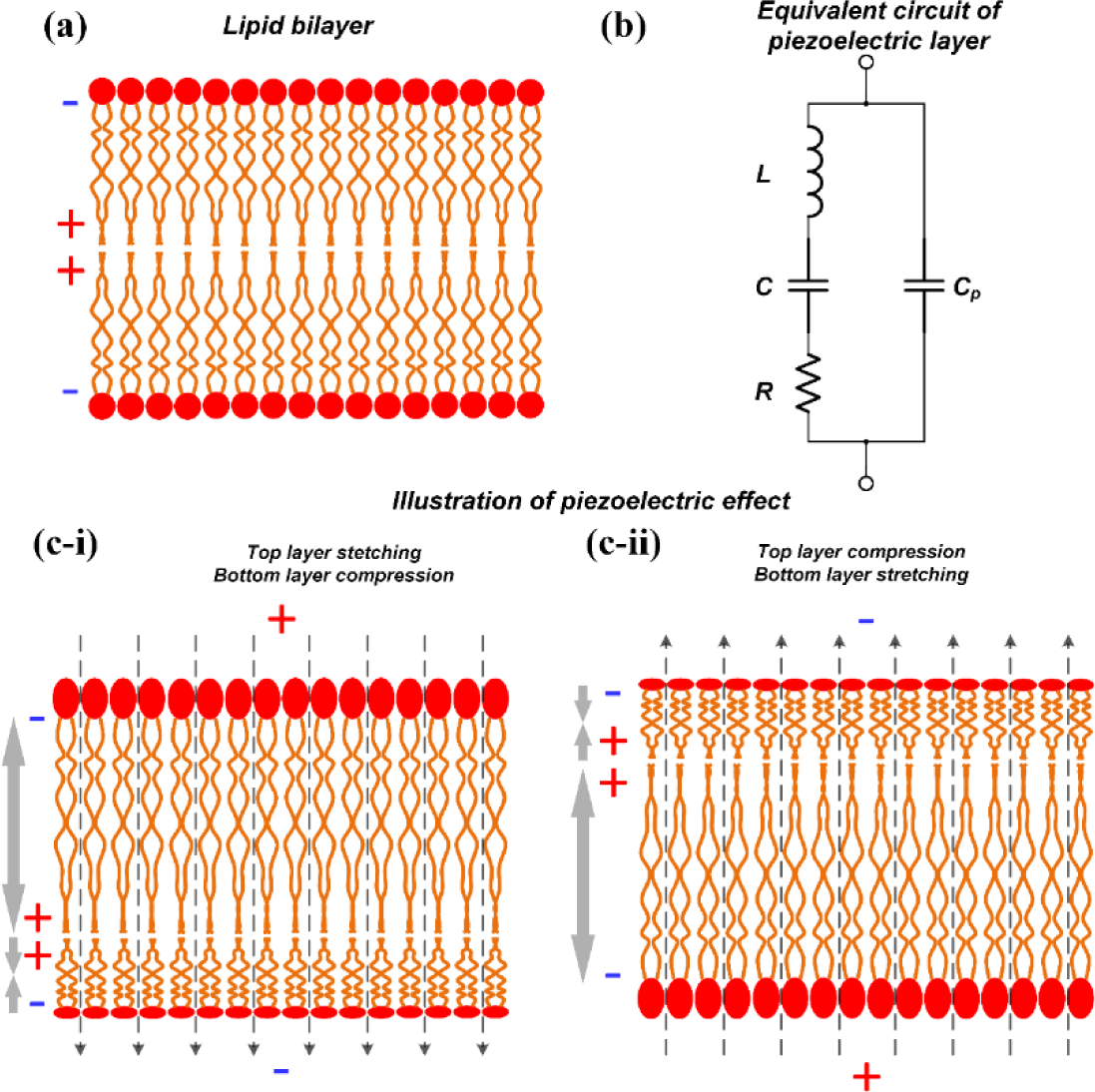
Illustration of the piezoelectric effect of the lipid bilayer structure of the cell membrane.

### B. Piezoelectric effect of plasma membrane

The plasma membrane is of a lipid bilayer structure as shown in Figure 13(a). The lipid molecules are polar molecules, with positive tails toward the center and negative tails toward the extra‐ and intracellular fluid (shown as the + and − signs) *(57)*. Then when an external electric field is applied onto this lipid bilayer membrane, the two layers will have opposite deformations as shown in Figure 13(c). When the electric field is top positive and bottom negative (shown in (c-i)), the top layer will be stretched and bottom layer will be compressed. When the electric field is top negative and bottom positive (shown in (c-ii)), the top layer will be compressed and bottom layer will be stretched. Here in the Figure 13(c), only axial deformation is depicted. However, the actual deformation will happen along all directions, resulting in a change of the surface tension. This surface tension can be measured not only in axons, but also in any cells with such lipid bilayer membranes *(58-60)*. On the other hand, a surface tension applied upon the lipid bilayer membrane can also change its electrostatic potential *(61)* and polarity *(62)*.

This phenomenon is called piezoelectric effect in MEMS (Microelectromechanical systems). The equivalent circuit of this piezoelectric layer is shown in Figure 13(b) *(63)*. In the right branch, ***C_P_*** is the capacitance of the cell membrane, determined by its area, thickness and dielectric constant. Then in the left branch, the three parameters, ***L***, ***C*** and ***R***, are just modelling parameters to fit the experiment result. They don’t have exact physical meanings. Here the value of ***C*** must be much smaller than ***C_P_***.

As can be seen, there will be an inductor in the equivalent circuit. Meanwhile, the circuit configuration is exactly the same as the simplified circuit shown in Figure 1. The value of the inductance is determined by the capacitance of the cell membrane ***C_P_*** and the resonance frequency ***f***. The resonance frequency is determined by the mechanical structure of the cell membrane. Considering the cell membrane is very thin and soft, it can be modeled as a very long, very thin and very soft beam, which has a very low mechanical resonance frequency. In our previous experiment on myelinated nerves, this resonance frequency is between 1k to 2k Hz. In Cole’s early experiment on giant squid axon, which is a non-myelinated, this resonance frequency is around 300 Hz. Now it is easy to comprehend why the resonance frequency is so low. With this very low resonance frequency, a huge inductor in the circuit is inevitable, since it is just an equivalent inductor to fit the experiment result. This equivalent inductor will not induce any magnetic field, but introduce a mechanical surface stress.

Then as an easy prediction, this mechanical stress can also be measured as a mechanical wave, which is accompany with the propagation of the action potential. This prediction was also made by Kenneth S. Cole at 1941 *(64)* after he measured the huge inductance in squid giant axon *(3)*. Even without any evidence at that time, Cole still believe it is the piezoelectric effect of plasma membrane to introduce this huge inductance. Since this mechanical wave is directly induced by the electric signal, it also has the same propagation speed as the electric signal of the action potential. This mechanical wave has been measured and studied for more than 10 years *(36)* and to be observed with not only exactly the same speed but also the same phase with the action potential *(65)*. An alternative theory, called soliton theory, was developed based on the observation of this mechanical wave. Based on the calculation of soliton theory, the propagation speed of the mechanical wave will be close to the action potential. However, after understanding the physical mechanism to generate this mechanical wave, we can immediately know this even without any calculation.

Moreover, this piezoelectric effect of the plasma membrane also explains the mechanism of acoustic nerve stimulation. Previous, there are several different possible mechanisms were proposed to explain why ultrasound can stimulate the nerve *(66)*. Now since the cell membrane is a piezoelectric film, the surface stress can be generated by applying electric field, and on the other hand, the surface stress can also generate electric field. The external applied acoustic wave can introduce this surface stress and then generate a voltage on the plasma membrane to activate the ion channels. This mechanism is totally the same as that of the electric stimulation. The only difference is that, this voltage on the plasma membrane is not induced by electrical coupling but by piezoelectric effect.

In summary, with the understanding of the inductance of myelin sheaths and the piezoelectric effect of the plasma membrane, a series of phenomena can be well explained. Meanwhile, a new explanation of the function of myelin can directly be obtained from this theory. Most importantly, it provides a comprehensive understanding of the nature of the neural signal: the neural signal is a combination of electric, magnetic and mechanical signals.

## Acknowledgments

We would like to thank for the experiment setup support from Han Wu, Shih Chiang Liu, Astrid, Shuhao Lu, Li Jing Ong and Dian Sheng Wong. We also would like to thank for the animal experiment support from Gammad Gil Gerald Lasam. We have our special acknowledgement to James T. Fulton for his pioneer research of neuroscience published on the Internet. This work was supported by grants from the National Research Foundation Competitive research programme (NRF-CRP) ‘Peripheral Nerve Prostheses: A Paradigm Shift in Restoring Dexterous Limb Function’ (NRF-CRP10-2012-01), National Research Foundation Competitive research programme (NRF-CRP) ‘Energy Harvesting Solutions for Biosensors’ (R-263-000-A27-281), National Research Foundation Competitive research programme (NRF-CRP) ‘Piezoelectric Photonics Using CMOS Compatible AlN Technology for Enabling The Next Generation Photonics ICs and Nanosensors’ (R-263-000-C24-281), Faculty Research Committee (FRC) ‘Thermoelectric Power Generator (TEG) Based Self-Powered ECG Plaster - System Integration (Part 3)’ (R-263-000-B56-112) and HIFES Seed Funding ‘Hybrid Integration of Flexible Power Source and Pressure Sensors’ (R-263-501-012-133). The theory was developed by Hao Wang, Jiahui Wang and Tianyiyi He. The modeling work was carried out by Hao Wang. The major framework of experiment design was carried out by Hao Wang, Jiahui Wang and Xin Yuan Thow. The experiments of CP nerve stimulation were carried out by Sanghoon Lee, Jiahui Wang, Xin Yuan Thow and Hao Wang. The experiments of TA muscle stimulation were carried out by Jiahui Wang, Xin Yuan Thow and Hao Wang. The experiments of cortical stimulation were carried out by Xin Yuan Thow, Jiahui Wang and Hao Wang. The experiments of pelvic stimulation were carried out Wendy Yen Xian Peh, Yuan Thow, Jiahui Wang and Hao Wang. The experiments of stimulus artifact recording with high sampling frequency system were carried out by Kian Ann Ng, Xin Yuan Thow, Jiahui Wang and Hao Wang. The data analysis was carried out by Hao Wang, Jiahui Wang, Xin Yuan Thow and Wendy Yen Xian Peh. The manuscript was written by Hao Wang, Jiahui Wang and Xin Yuan Thow. All authors discussed the experimental results and contributed to the final version of the manuscript. Prof. Nitish V. Thakor, Prof. Chengkuo Lee and Prof. Chia-hung Chen provided general guidance and supervision of the project. Authors declare no competing interests. All data is available in the main text or the supplementary materials.

## Supplementary Materials

### Method

S1. Cortical stimulation with sciatic nerve recording test electrode configuration, testing setup, detailed procedure and testing parameters

S2. Pelvic nerve test electrode configuration, testing setup, detailed procedure and testing parameters

Revised circuit

S3. Circuit with additional capacitor Additional data:

S4 Repeated data for stimulus artifacts Other theoretical analysis:

S5 The answer to the two questions S6 Circuit analysis

## References and Notes

(1) Brzychczy, S. and Poznanski, R.R., 2013. Mathematical neuroscience. Academic Press.

(2) Hodgkin, A.L. and Huxley, A.F., 1952. A quantitative description of membrane current and its application to conduction and excitation in nerve. The Journal of physiology, 117(4), pp.500-544.

(3) Cole, K.S. and Baker, R.F., 1941. Longitudinal impedance of the squid giant axon. The Journal of general physiology, 24(6), pp.771-788.

(4) Rossi, S. and Griffith, B.E., 2017. Incorporating inductances in tissue-scale models of cardiac electrophysiology. Chaos: An Interdisciplinary Journal of Nonlinear Science, 27(9), p.093926.

(5) Scott, A.C., 1971. Effect of the series inductance of a nerve axon upon its conduction velocity. Mathematical Biosciences, 11(3-4), pp.277-290.

(6) Mosgaard, L.D., Zecchi, K.A., Heimburg, T. and Budvytyte, R., 2015. The effect of the nonlinearity of the response of lipid membranes to voltage perturbations on the interpretation of their electrical properties. A new theoretical description. Membranes, 5(4), pp.495-512.

(7) Hutcheon, B. and Yarom, Y., 2000. Resonance, oscillation and the intrinsic frequency preferences of neurons. Trends in neurosciences, 23(5), pp.216-222.

(8) Dwyer, J., Lee, H., Martell, A. and van Drongelen, W., 2012. Resonance in neocortical neurons and networks. European Journal of Neuroscience, 36(12), pp.3698-3708.

(9) Araki, T., Ito, M. and Oshima, T., 1961. Potential changes produced by application of current steps in motoneurones. Nature, 191(4793), p.1104.

(10) Ranck Jr, J.B., 1963. Analysis of specific impedance of rabbit cerebral cortex. Experimental neurology, 7(2), pp.153-174.

(11) Freeman, W.J., 1961. Harmonic oscillation as model for cortical excitability changes with attention in cats. Science, 133(3470), pp.2058-2059.

(12) Homble, F. and Jenard, A., 1984. Pseudo-inductive behaviour of the membrane potential of Chara corallina under galvanostatic conditions: a time-variant conductance property of potassium channels. Journal of experimental botany, 35(9), pp.1309-1322.

(13) Huxley, A.F., 1963. The quantitative analysis of excitation and conduction in nerve. Les Prix Nobel, 1963, pp.242-260.

(14) Guttman, R., 1969. Temperature dependence of oscillation in squid axons: comparison of experiments with computations. Biophysical journal, 9(3), p.269.

(15) Sjodin, R.A. and Mullins, L.J., 1958. Oscillatory behavior of the squid axon membrane potential. The Journal of general physiology, 42(1), pp.39-47.

(16) Curtis, H.J. and Cole, K.S., 1942. Membrane resting and action potentials from the squid giant axon. Journal of cellular and comparative physiology, 19(2), pp.135-144.

(17) Thomas, A., 2013. Memristor-based neural networks. Journal of Physics D: Applied Physics, 46(9), p.093001.

(18) Mauro, A., Conti, F., Dodge, F. and Schor, R., 1970. Subthreshold behavior and phenomenological impedance of the squid giant axon. The Journal of general physiology, 55(4), pp.497-523.

(19) Howell, B., Medina, L.E. and Grill, W.M., 2015. Effects of frequency-dependent membrane capacitance on neural excitability. Journal of neural engineering, 12(5), p.056015.

(20) Li, C.L. and Bak, A., 1976. Excitability characteristics of the A-and C-fibers in a peripheral nerve. Experimental neurology, 50(1), pp.67-79.

(21) Evans, E.F., 1972. The frequency response and other properties of single fibres in the guinea pig cochlear nerve. The Journal of physiology, 226(1), pp.263-287.

(22) Kral, A., Hartmann, R., Mortazavi, D. and Klinke, R., 1998. Spatial resolution of cochlear implants: the electrical field and excitation of auditory afferents. Hearing research, 121(1-2), pp.11-28.

(23) Hartmann, R., Topp, G. and Klinke, R., 1984. Discharge patterns of cat primary auditory fibers with electrical stimulation of the cochlea. Hearing research, 13(1), pp.47-62.

(24) Frankenhaeuser, B. and Widén, L., 1956. Anode break excitation in desheathed frog nerve. The Journal of physiology, 131(1), pp.243-247.

(25) Wright, E.B. and Ooyama, H., 1961. Anode break excitation and Pflüger’s law. American Journal of Physiology-Legacy Content, 200(2), pp.219-222.

(26) Howell, B., Medina, L.E. and Grill, W.M., 2015. Effects of frequency-dependent membrane capacitance on neural excitability. Journal of neural engineering, 12(5), p.056015.

(27) Ranjan, R., Chiamvimonvat, N., Thakor, N.V., Tomaselli, G.F. and Marban, E., 1998. Mechanism of anode break stimulation in the heart. Biophysical journal, 74(4), pp.1850-1863.

(28) Weiss, G., 1990. Sur la possibilite de rendre comparables entre eux les appareils servant a l’excitation electrique. Archives Italiennes de Biologie, 35(1), pp.413-445.

(29) Lapicque, L., 1909. Definition experimentale de 1’excitabilite. Soc Biol., 77, pp.280-283.

(30) Su, X., Simenson, H.A., Dinsmoor, D.A. and Orser, H.D., 2017. Evaluation of Pulse-Width of Spinal Nerve Stimulation in a Rat Model of Bladder Micturition Reflex. Neuromodulation: Technology at the Neural Interface, 20(8), pp.793-798.

(31) Friedli, W.G. and Meyer, M., 1984. Strength-duration curve: a measure for assessing sensory deficit in peripheral neuropathy. Journal of Neurology, Neurosurgery & Psychiatry, 47(2), pp.184-189.

(32) Rodríguez-Fernández, Á.L., Rebollo-Roldán, J., Jiménez-Rejano, J.J. and Güeita-Rodríguez, J., 2016. Strength-duration curves of the common fibular nerve show hypoexcitability in people with functional ankle instability. PM&R, 8(6), pp.536-544.

(33) Uzman, B.G. and Nogueira-Graf, G., 1957. Electron microscope studies of the formation of nodes of Ranvier in mouse sciatic nerves. The Journal of Cell Biology, 3(4), pp.589-598.

(34) Armati, P.J. and Mathey, E.K., 2013. An update on Schwann cell biology— immunomodulation, neural regulation and other surprises. Journal of the neurological sciences, 333(1-2), pp.68-72.

(35) Bunge, R.P., Bunge, M.B. and Bates, M., 1989. Movements of the Schwann cell nucleus implicate progression of the inner (axon-related) Schwann cell process during myelination. The Journal of Cell Biology, 109(1), pp.273-284.

(36) Appali, R., van Rienen, U. and Heimburg, T., 2012. A comparison of the Hodgkin–Huxley model and the soliton theory for the action potential in nerves. In Advances in Planar Lipid Bilayers and Liposomes (Vol. 16, pp. 275-299). Academic Press.

(37) Legatt, A.D., Schroeder, C.E., Gill, B. and Goodrich, J.T., 1992. Electrical stimulation and multichannel EMG recording for identification of functional neural tissue during cauda equina surgery. Child’s Nervous System, 8(4), pp.185-189.

(38) Frigo, C., Ferrarin, M., Frasson, W., Pavan, E. and Thorsen, R., 2000. EMG signals detection and processing for on-line control of functional electrical stimulation. Journal of Electromyography and Kinesiology, 10(5), pp.351-360.

(39) Lieberstein, H.M. and Mahrous, M.A., 1970. A source of large inductance and concentrated moving magnetic fields on axons. Mathematical Biosciences, 7(1-2), pp.41-60.

(40) Seipel, J.H. and Morrow, R.D., 1960. The Magnetic Field Accompanying Neuronal Activity A New Method for The Study of The Nervous System. Journal of the Washington Academy of Sciences, 50(6), pp.1-4.

(41) Gengerelli, J.A., Holter, N.J. and Glasscock, W.R., 1961. Magnetic fields accompanying transmission of nerve impulses in the frog’s sciatic. The Journal of Psychology, 52(2), pp.317-326.

(42) Cohen, D., 1967. Magnetic fields around the torso: production by electrical activity of the human heart. Science, 156(3775), pp.652-654.

(43) Cohen, D., 1968. Magnetoencephalography: evidence of magnetic fields produced by alpha rhythm currents. Science, 161(3843), pp.784-786.

(44) Wikswo, J.P., Samson, P.C. and Giffard, R.P., 1983. A low-noise low input impedance amplifier for magnetic measurements of nerve action currents. IEEE Transactions on Biomedical Engineering, (4), pp.215-221.

(45) Wikswo, J.P., Barach, J.P. and Freeman, J.A., 1980. Magnetic field of a nerve impulse: first measurements. Science, 208(4439), pp.53-55.

(46) Barach, J.P., Freeman, J.A. and Wikswo Jr, J.P., 1980. Experiments on the magnetic field of nerve action potentials. Journal of Applied Physics, 51(8), pp.4532-4538.

(47) Wikswo Jr, J.P., 1981. Recent developments in the measurement of magnetic fields from isolated nerves and muscles. Journal of Applied Physics, 52(3), pp.2554-2559.

(48) Wikswo Jr, J.P., 1982. Improved instrumentation for measuring the magnetic field of cellular action currents. Review of Scientific Instruments, 53(12), pp.1846-1850.

(49) Roth, B.J. and Wikswo, J.P., 1985. The magnetic field of a single axon. A comparison of theory and experiment. Biophysical journal, 48(1), pp.93-109.

(50) Barach, J.P., Roth, B.J. and Wikswo, J.P., 1985. Magnetic measurements of action currents in a single nerve axon: A core-conductor model. IEEE Trans Biomed Eng, 32(2), pp.136-140.

(51) Wijesinghe, R.S., 2010. Magnetic measurements of peripheral nerve function using a neuromagnetic current probe. Experimental Biology and Medicine, 235(2), pp.159-169.

(52) Wikswo, J.P., Barach, J.P., Gundersen, S.C., McLean, M.J. and Freeman, J.A., 1983. First magnetic measurements of action currents in isolated cardiac Purkinje fibers. Il Nuovo Cimento D, 2(2), pp.368-378.

(53) Hallett, M., 2007. Transcranial magnetic stimulation: a primer. Neuron, 55(2), pp.187-199.

(54) Maccabee, P.J., Amassian, V.E., Eberle, L.P. and Cracco, R.Q., 1993. Magnetic coil stimulation of straight and bent amphibian and mammalian peripheral nerve in vitro: locus of excitation. The Journal of Physiology, 460(1), pp.201-219.

(55) Davids, M., Guérin, B., Malzacher, M., Schad, L.R. and Wald, L.L., 2017. Predicting Magnetostimulation Thresholds in the Peripheral Nervous System using Realistic Body Models. Scientific reports, 7(1), p.5316.

(56) Irnich, W., 1994. Electrostimulation by time-varying magnetic fields. Magnetic Resonance Materials in Physics, Biology and Medicine, 2(1), pp.43-49.

(57) Yang, Y., Mayer, K.M., Wickremasinghe, N.S. and Hafner, J.H., 2008. Probing the lipid membrane dipole potential by atomic force microscopy. Biophysical journal, 95(11), pp.5193-5199.

(58) Chiu, S.W., Clark, M., Balaji, V., Subramaniam, S., Scott, H.L. and Jakobsson, E., 1995. Incorporation of surface tension into molecular dynamics simulation of an interface: a fluid phase lipid bilayer membrane. Biophysical journal, 69(4), pp.1230-1245.

(59) Brockman, H., 1994. Dipole potential of lipid membranes. Chemistry and physics of lipids, 73(1-2), pp.57-79.

(60) Feller, S.E., 2000. Molecular dynamics simulations of lipid bilayers. Current opinion in colloid & interface science, 5(3-4), pp.217-223.

(61) Warshaviak, D.T., Muellner, M.J. and Chachisvilis, M., 2011. Effect of membrane tension on the electric field and dipole potential of lipid bilayer membrane. Biochimica et Biophysica Acta (BBA)-Biomembranes, 1808(10), pp.2608-2617.

(62) Zhang, Y.L., Frangos, J.A. and Chachisvilis, M., 2006. Laurdan fluorescence senses mechanical strain in the lipid bilayer membrane. Biochemical and biophysical research communications, 347(3), pp.838-841.

(63) Richter, B., Twiefel, J. and Wallaschek, J., 2009. Piezoelectric equivalent circuit models. In Energy Harvesting Technologies(pp. 107-128). Springer, Boston, MA.

(64) Cole, K.S., 1941. Rectification and inductance in the squid giant axon. The Journal of general physiology, 25(1), pp.29-51.

(65) Gonzalez-Perez, A., Mosgaard, L.D., Budvytyte, R., Villagran-Vargas, E., Jackson, A.D. and Heimburg, T., 2016. Solitary electromechanical pulses in lobster neurons. Biophysical chemistry, 216, pp.51-59.

(66) Wright, C.J., Rothwell, J. and Saffari, N., 2015. Ultrasonic stimulation of peripheral nervous tissue: an investigation into mechanisms. In Journal of Physics: Conference Series (Vol. 581, No. 1, p. 012003). IOP Publishing.

